# Enhancing Ebola Virus Disease Surveillance And Prevention In Counties Without Confirmed Cases In Rural Liberia: Experiences From Sinoe County During The Flare-Up In Monrovia, April To June, 2016

**DOI:** 10.1101/139154

**Authors:** Vera Darling Weah, John S. Doedeh, Samson Q. Wiah, Emmanuel Nyema, Siafa Lombeh, Jeremias Naiene

## Abstract

**Introduction:** During the flare-ups of Ebola virus disease (EVD) in Liberia, Sinoe County reactivated the multi-sectorial EVD control strategy in order to be ready to respond to the eventual reintroduction of cases.

This paper describes the impacts of the interventions implemented in Sinoe County during the last flare-up in Monrovia, from April 1 to June 9, 2016, using the resources provided during the original outbreak that ended one year back.

**Methods:** We conducted a descriptive study to describe the key interventions implemented in Sinoe County, the capacity available, the implications for the reactivation of the multi-sectoral EVD control strategy, and the results of the same. We also conducted a cross-sectional study to analyze the impact of the interventions on the surveillance and on infection prevention and control (IPC).

**Results:** The attrition of the staff trained during the original outbreak was low, and most of the supplies, equipment, and infrastructure from the original outbreak remained available.

With an additional US$1755, improvements were observed in the IPC indicators of triage, which increased from a mean of 60% during the first assessment to 77% (P=0.002). Additionally, personal/staff training improved from 78% to 89% (P=0.04).

The percentage of EVD death alerts per expected deaths investigated increased from 26% to 63% (P<0.0001).

**Discussion:** The small attrition of the trained staff and the availability of most of the supplies, equipment, and infrastructure made the reactivation of the multi-sectoral EVD control strategy fast and affordable. The improvement of the EVD surveillance was possibly affected by the community engagement activities, awareness and mentoring of the health workers, and improved availability of clinicians in the facilities during the flare-up. The community engagement may contribute to the report of community-based events, specifically community deaths. The mentoring of the staff during the supportive supervisions also contributed to improve the IPC indicators.

## Introduction

The Ebola virus disease (EVD) outbreak started in Guinea in 2013 (1,2), and as of June 10, 2016, 28616 cases had been registered, with 11310 deaths (3). After the end of the original outbreak in the three most affected countries in 2015 (2), specifically, in Liberia in May, in Sierra Leone in November, and in Guinea in December, different flare-ups were reported. The biggest flare-up was in Guinea, which occurred from February 27 (4) to June 1, 2016, with 10 reported cases and seven deaths (3), while the smallest one was in Sierra Leone, from January 14 to March 17, 2016, with two reported cases and one death (5).

The flare-ups may occur due to importation, reintroduction of the virus from animal reservoir, missed chain of transmission, and reemergence of virus from a survivor (4,6–10), and can be easily detected when EVD surveillance, including the community-based surveillance and laboratory capacity, is established (11). EVD flare-up can also be controlled on time when a multi-sectorial EVD control strategy is implemented effectively. This strategy involves different committees, including clinical case management, surveillance, laboratory, logistic, behavioral and social interventions, psychosocial support, coordination, and others (11).

Liberia reported three flare-ups after the initial declaration of “disease free” status on May 9, 2015 (2,12,13), the first one being from June 29 to September 3, 2015 in Margibi County (14), which occurred after the re-emergence of the virus from a survivor through sexual contact (12,14), and the second one being in Duport road, Monrovia, from November 24, 2015 (15,16) to January 14, 2016, which started from a pregnant Ebola survivor who became infectious when her immune system weakened due to the pregnancy (15). The two flare-ups were detected through a postmortem swab tested for EVD (14,15). The last flare-up, which occurred from April 1 to June 9, 2016, was imported from Guinea (3).

All the flare-ups were detected early, during the 90 days of enhanced EVD surveillance recommended after the end of the outbreak, which includes the swabbing of all the dead bodies for EVD laboratory investigation (17). The active Incident Management System (IMS) for coordination, a temporary field based Emergency Operation Center (EOC), implementation of the rapid response plan developed to respond to eventual flare-ups, and the presence of experienced staff trained during the original outbreak contributed to early containment of the flare-ups (14).

Sinoe County reported 22 confirmed EVD cases and 11 deaths during the original outbreak that was controlled with the isolation of the cases (18), establishment of Ebola task forces, training of the staff, and other strategies (19,20). The last confirmed case died in December 2014 and no flare-up was reported in the county. However, during the flare-ups in Liberia and neighboring countries, Sinoe County reactivated the multi-sectorial EVD control strategy in order to be ready to respond to the eventual reintroduction of cases. This paper describes the impacts of the interventions implemented in Sinoe County during the last flare-up in Monrovia, using the resources provided during the original outbreak, in order to be ready to respond to the eventual importation of cases.

## Methods

### Setting

Sinoe County, one of the southeastern counties in rural Liberia, is divided into 10 health districts, four of which have a history of EVD positive cases reported during the original outbreak (Fig 1) (21). The capital city Greenville is located at about 150 miles from the capital of Liberia, Monrovia. The population is dispersed, with 104,932 inhabitants and a density of 27 people per square mile (22). It is difficult to reach many communities owing to forests, rivers, swamps, and hills, and the average distance from communities to healthcare facilities is 6.6 km (21,23). The county is served by two medical doctors, 18 physician assistants, and 67 nurses, and it has 35 health facilities, including one referral hospital with a capacity of 100 beds, and 34 clinics (21).

**Fig 1:**
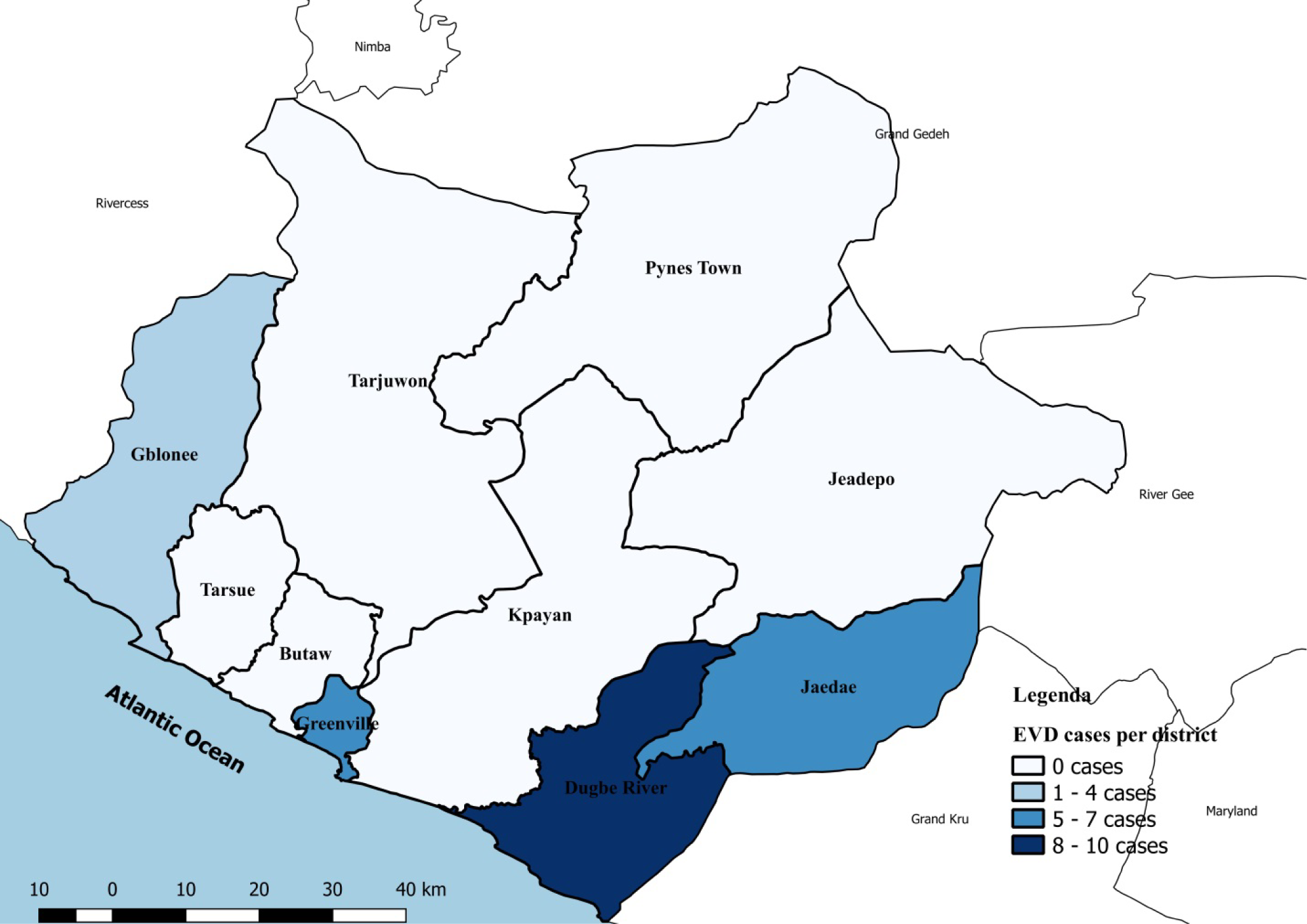
Map of Sinoe County, Liberia showing the number of EVD cases reported per district during the original outbreak, 2014.

### Study design

We conducted a descriptive study to describe the key interventions implemented in the county from April 1 to June 9, 2016, the capacity available, the implications for the reactivation of the multi-sectoral EVD control strategy, and the results of the same. We also conducted a cross-sectional study to analyze the impact of the interventions on surveillance and infection, prevention and control (IPC).

### Data analysis

We entered the data into Microsoft Excel^™^ and used MedCalc^®^ Statistical Software version 17.2 (24) for IPC and EVD surveillance data analysis. To determine statistical significance, we performed the students-t test for the IPC assessment data and the Chi-squared test for the EVD surveillance data. We calculated the number of expected deaths using the crude death rate in Liberia, of 8.8 deaths/1000 population/year (25,26).

### Key interventions

We reactivated the different committees involved in the EVD control activities, as recommended by the WHO (Fig 2) (11).

**Fig 2:**
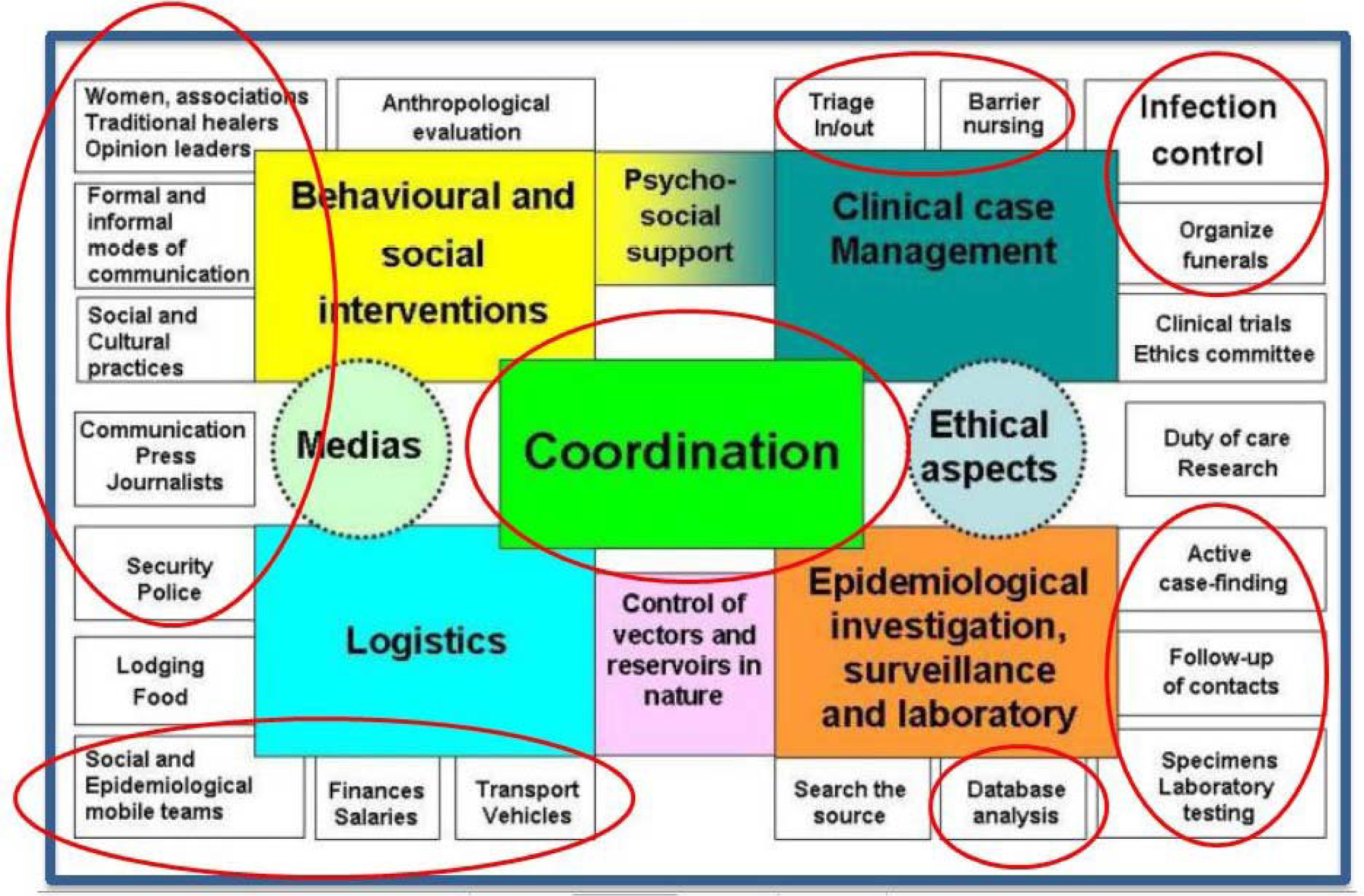
Organizational structures of the EVD control activities (11) highlighting the committee activated in Sinoe County, Liberia during the flare-up, 2016.

### Coordination

The county health team (CHT) developed an emergency plan to prepare for responding to the eventual importation of cases from Monrovia using the resources provided during the original outbreak. When the end of the flare-up was declared, we analyzed the level of implementation of the plan as well as the costs involved.

The coordination committee conducted meetings with county, district, and community stakeholders, partners, and other line ministries for coordination, awareness, and advocacy, in order to mobilize resources.

### Logistics and human resources

We analyzed the human resources database to assess how many Rapid Response Team (RRT) members, contact tracers, and burial team members trained during the original outbreak were available in the county during the last flare-up.

Using the minimum standards assessment tool developed during the original outbreak we conducted the first integrated assessment of the logistic capacity available at 30 (88%) health facilities in the county, including the referral hospital, in April 2016. However, five (14%) of the 35 health facilities in county were inaccessible.

After the initial assessment we replenished the supplies, mentored the health care workers, and conducted the second assessment from the end of May to October, 2016 to verify the changes. The assessments were conducted through direct observation, interviews of the healthcare workers, and perusal of the documents available.

We also assessed the 10 most important check points connecting Sinoe County and other counties to verify the knowledge of the staff and availability of IPC supplies.

### Clinical case investigation, surveillance, and laboratory

We conducted supportive supervisions at 30 (88%) health facilities in Sinoe County and held weekly meetings with district health officers and district surveillance officers to reinforce the triage of all the patients, the use of the EVD outbreak case definitions, and to analyze the EVD surveillance situation in the county.

Besides using the WHO case definition for the investigation of cases (Figure 3) (27) we collected swabs of all the dead bodies, regardless of the cause of death, to perform *Real*-*time* quantitative reverse transcription *PCR* (qRT-*PCR*) before and during the flare-up, as part of the 90 days of enhanced surveillance implemented after the end of each EVD outbreak.

The specimens were collected by trained clinicians and lab staff in all the health facilities in the county and were transported to the regional labs in Liberia with the capacity to perform the qRT-PCR for EVD.

We perused all the lab records to quantify the numbers of specimens collected before and during the flare-up.

**Figure 3:** World Health Organization case definition of EVD used during the outbreaks and used in Sinoe County before and during the flare-up in Liberia, from April to June 2016.

**SUSPECTED CASE:**

Any person, alive or dead, suffering or having suffered from a sudden onset of high fever and having had contact with

- a suspected, probable or confirmed Ebola or Marburg case
- a dead or sick animal (for Ebola)

**OR:**

Any person with sudden onset of high fever and at least three of the following symptoms:

- headaches
- vomiting
- anorexia / loss of appetite
- diarrhea
- lethargy
- stomach pain
- aching muscles or joints
- difficulty swallowing
- breathing difficulties
- hiccup

**OR:**

Any person with inexplicable bleeding

### Behavioral and social interventions

We disseminated EVD prevention messages through radio talk shows with county authorities and traditional leaders at a local radio station. We also conducted community meetings in high risk communities, churches, mosques, and funeral and healing homes to increase awareness and to encourage the reporting of community deaths to the health facilities.

### Ethical considerations

Ethical approval was not required to implement the activities since they were part of the activities of the Ministry of Health to respond to outbreaks in Liberia. We did not use any confidential data and did not disclose any unauthorized names in our report.

## Results

### Coordination

The county’s task force for EVD was activated and was responsible to ensure that the preventive measures were implemented in all levels and that any suspected case was promptly reported. Since the resources were already available from the original outbreak, the three-month plan costed an additional US$1755 besides the budget for the routine activities. The dissemination of EVD prevention messages to churches, mosques, households, meetings in high risk communities, and funeral and healing homes was the most expensive activity, and it costed about US$400, especially to purchase fuel for the activities and to pay daily subsistence allowance (DSA) for the staff (Table 1).

**Table 1:**
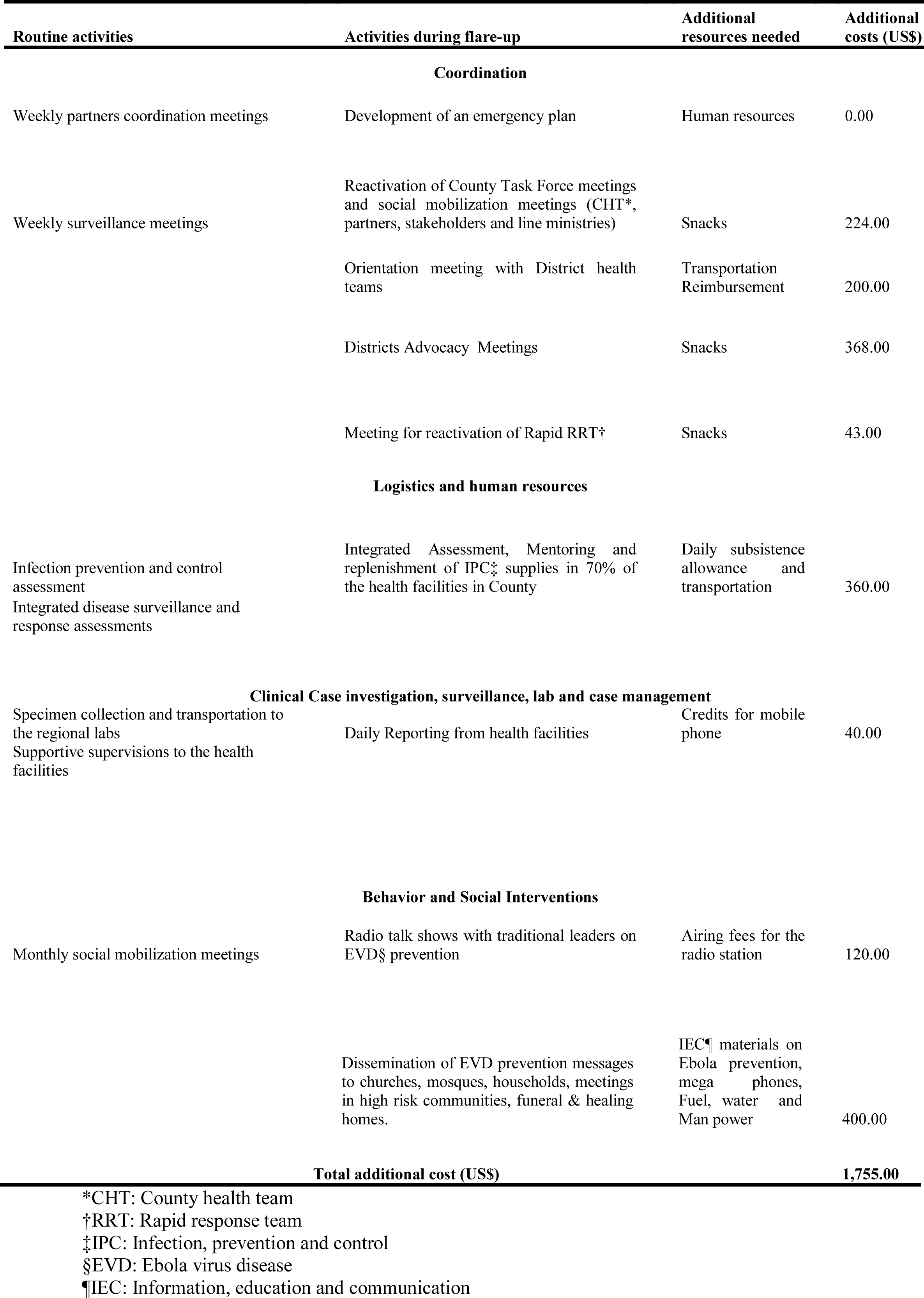
List of the main activities implemented routinely and during the flare-up for EVD prevention and surveillance in Sinoe County, Liberia 2016

### `Logistics, human resources and, Infection Prevention and Control assessments

At the first assessment, we included 30 health facilities (29 clinics and one hospital) in our analysis of the IPC indicators. From this analysis, we excluded the indicators that were not applicable to clinics (Appendix 1), according to the tool used. Personal/staff training was the group of indicators with higher score. The mean of the four individual indicators in this group was 78% [standard deviation (SD)=11] including 90% of the health facilities with staffs trained in IPC by the Ministry of Health and Social Welfare (MOHSW) in a training called “keep safe keep serving.” Additionally, 63% with staffs meeting the criteria outlined in the MOHSW’s Essential Package of Health Services (EPHS), the minimum skills required to work in the facilities (Table 2). On the other hand, the group of four indicators for triage had the mean of 60% (SD=12%) and the other three indicators assessing the facilities in terms of having an appropriate isolation space ready to receive cases had the mean of 52% (SD=9%).

**Table 2:**
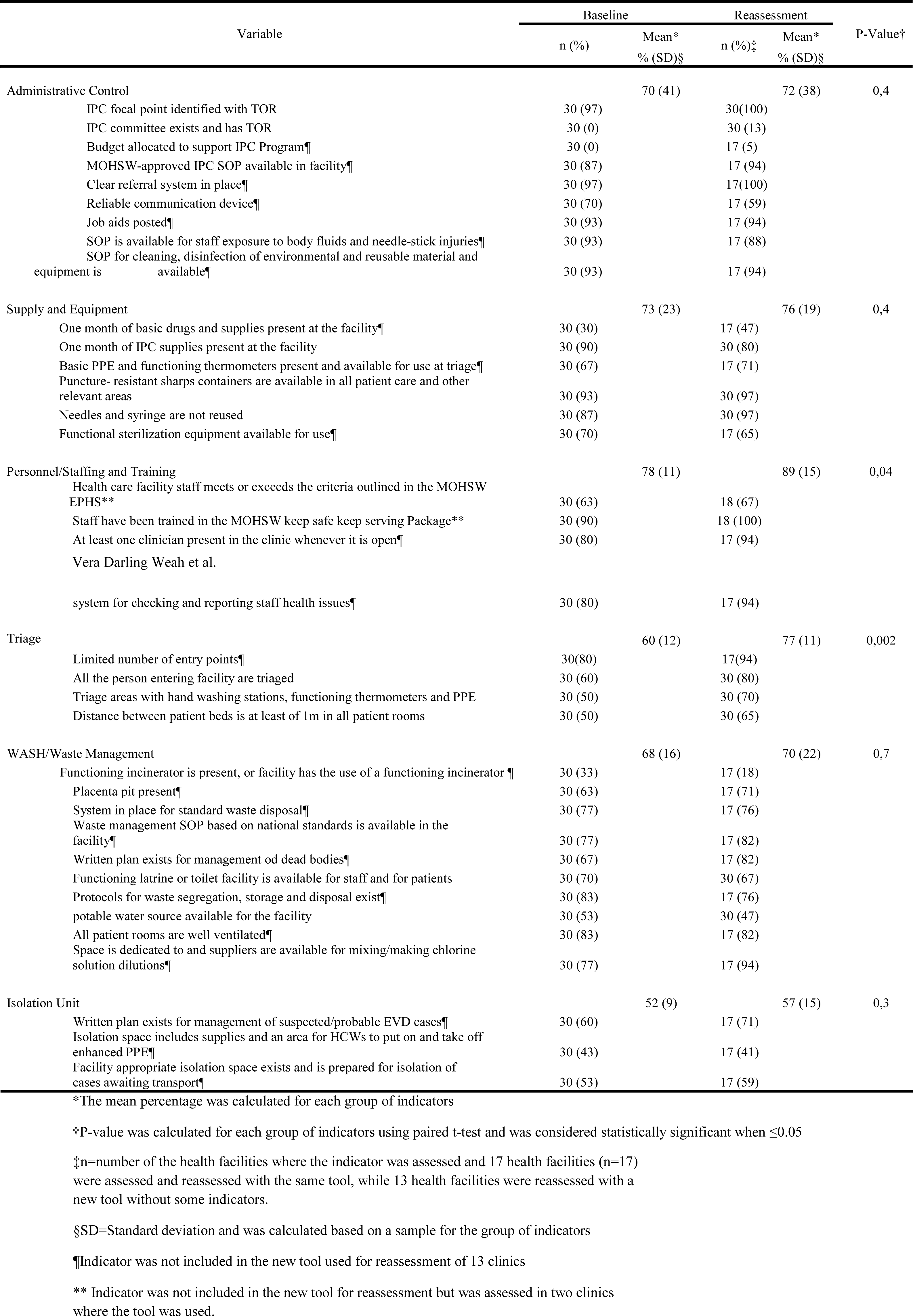
Performance of the IPC indicators in Sinoe County at the beginning (baseline) and at the end (reassessment) of the flare-up in Monrovia, Liberia 2016

The structure of the community care center in the Karquekpo community, with a capacity of 12 beds, and the ETU in Greenville district, with a capacity of 60 beds, remained intact, but these facilities required equipment and supplies to start receiving patients.

Out of the facilities assessed, 27 (90%) had IPC supplies that would last them for one month. These supplies were provided during the original outbreak.

In the second assessment, significant improvements were observed on the indicators of triage (an increase from 60% during the first assessment to 77% on reassessment; P=0.002) and personal/staff training (an increase from 78% to 89%; P=0.04), while other indicators did not exhibit significant improvements.

The lab supplies for the oral swab of dead bodies and whole blood for live alert investigation were available in all the facilities (100%) during the first and the second assessment.

The county had three vehicles that were available to be used in case of an outbreak, including one ambulance for the transportation of cases.

From the 21 members of the RRT trained during the original outbreak, 19 (90.5%) remained present in the county. All the 250 (100%) community volunteers trained for contact tracing remained in their communities and they were available to resume the task in case of need. From the the 14 members of the two burial teams trained during the previous outbreak, 12 (86%) remained present in the county.

All the 10 most important check points connecting Sinoe with the other counties had a bucket for handwashing and thermometers, although some were not functional. After the mentoring of the staff and replacement of the thermometers, nine (90%) check points reactivated the monitoring of temperature and handwashing for people crossing these locations.

### Clinical case investigation, surveillance, and laboratory

The percentage of EVD death alerts investigated, including the oral swabs collected and sent to the regional lab in Liberia, increased from 26% of the death alerts per expected deaths during the three months before the flare-up in Monrovia, to 63% of the alerts per expected deaths during the flare-up (P<0.0001). Significant improvement was verified in seven of the 10 health districts in the county. On the other hand, the number of live alerts investigated and the whole blood tested for EVD decreased from 0.5 alerts per 100 population to 0.4 alerts per 100 population (P=0.0003). The reduction was significant in Greenville (P<0.0001) and Butaw (P=0.05) districts (Table 3). All the specimens from both live and death alerts came negative for EVD.

The number of health facilities investigating death alerts increased from 19 (Mean=54% per health district, SD=34%) to 28 (Mean=84% per health district, SD=22%) health facilities (P=0.006) in all the 10 health districts in the county (Figure 4).

**Table 3:** Live and death alerts investigated in Sinoe County before and during the flare up in Monrovia, Liberia, 2016

| District | Population | Expected deaths<br>(12 weeks) | Live alerts |  |  | Death alerts |  |  |
| --- | --- | --- | --- | --- | --- | --- | --- | --- |
|  |  |  | Before the flare-up<br>January to March | During flare-up<br>April to June | P-Value* | Before the flare-up<br>January to March | During flare-up<br>April to June | P-Value* |
|  |  |  | Alerts (Alerts per<br>100 population) | Alerts (Alerts per<br>100 population) |  | Alerts (% of alerts per<br>expected deaths) | Alerts (% of alerts per<br>expected deaths) |  |
| Butaw | 5372 | 12 | 43 (0.8) | 26 (0.5) | 0.05 | 1(8) | 9(75) | 0.001 |
| Kpanyan | 13479 | 26 | 72 (0.5) | 70 (0.5) | 1 | 7(27) | 21(81) | 0.0001 |
| Tarjuowon | 24031 | 48 | 90 (0.4) | 62(0.3) | 0.06 | 1(2) | 24(50) | <0.0001 |
| Greenville | 18975 | 38 | 133 (0.7) | 65 (0.3) | <0.0001 | 26(68) | 36(95) | 0.003 |
| Dugbe river | 13377 | 26 | 49 (0.4) | 51 (0.4) | 1 | 19(73) | 18(69) | 0.8 |
| Gblonee | 5196 | 10 | 22 (0.4) | 16 (0.3) | 0.4 | 1(10) | 5(50) | 0.06 |
| Tarsue | 7648 | 16 | 26 (0.3) | 14 (0.2) | 0.2 | 0(0) | 9(56) | 0.0005 |
| Jeadae | 12154 | 24 | 31 (0.3) | 28 (0.2) | 0.1 | 4(17) | 17(71) | 0.0002 |
| Pynes | 10116 | 20 | 47 (0.5) | 64 (0.6) | 0.3 | 1(5) | 6(30) | 0.04 |
| Jedeapo | 7144 | 14 | 24 (0.3) | 30 (0.4) | 0.3 | 2(14) | 2(14) | 1 |
| **Total** | **117492** | **234** | **537 (0.5)** | **426 (0.4)** | **0.0003** | **62(26)** | **147(63)** | **<0.0001** |

\*P-value was calculated using Chi-squared test, comparison of two proportions and was considered statistically significant when $\leq 0.05$

**Figure 4:**
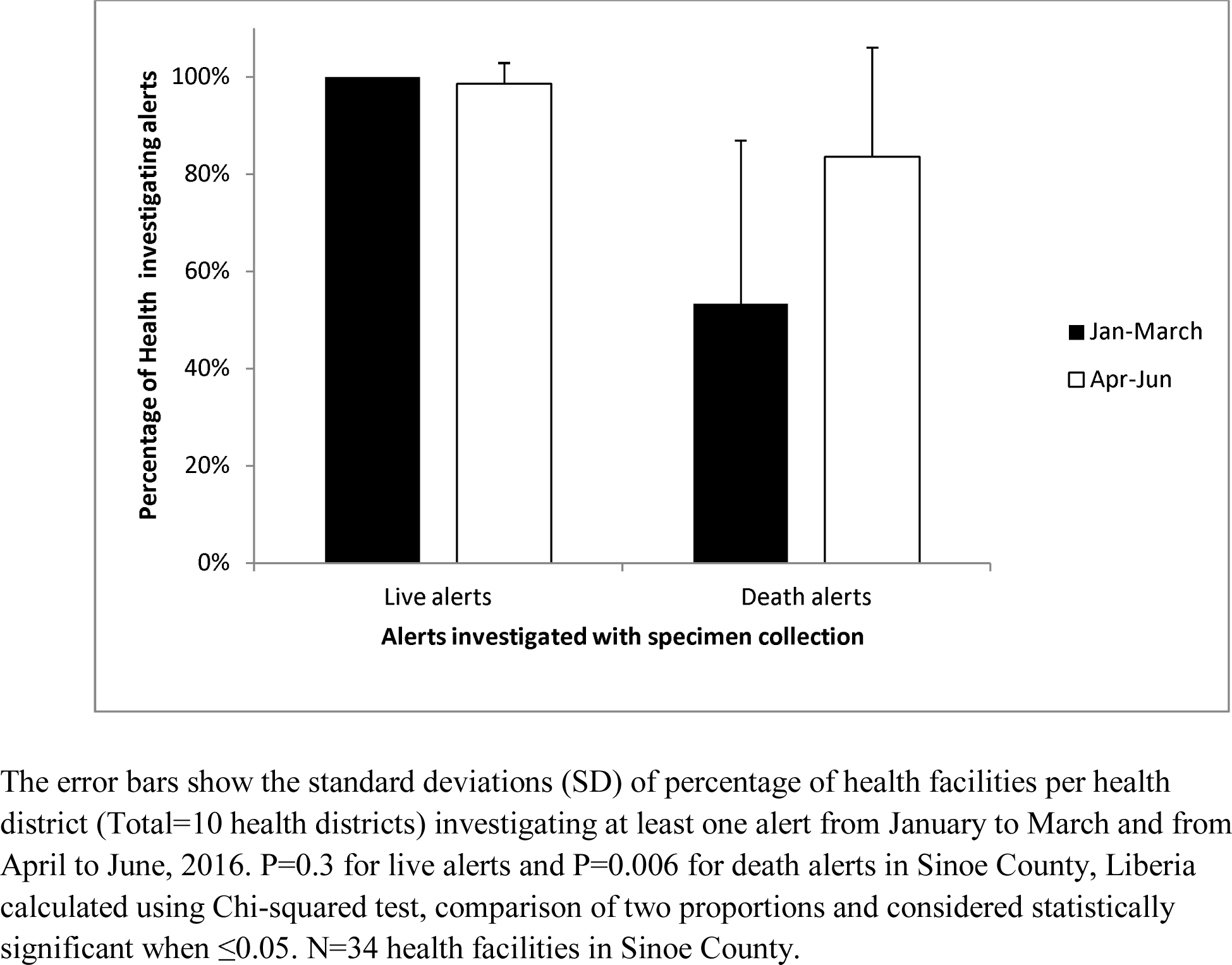
Graph of the percentage of health facilities investigating EVD alerts in Sinoe County three months before and three months during the flare-up in Monrovia, Liberia 2016

## Discussion

The interventions implemented in Sinoe County in preparation for responding to the reintroduction of EVD cases from the last flare-up in Monrovia had a significant impact on EVD surveillance, leading to an increase in percentage of death alerts from 26% alerts per expected deaths during the three months before, to 63% alerts per expected deaths during the flare-up (P<0.0001). This finding was possibly affected by the community engagement activities, awareness and mentoring of the health workers, and improved availability of clinicians in the facilities during the flare-up. Similar findings were reported in Lofa County, Liberia, during the original outbreak in 2014, when more community deaths were reported to the health authorities to be investigated for EVD after community sensitization and acceptance (28). The reasons for reduction of live alerts were not clear from our study, and the same may have occurred owing to the increased attention focused on the death alerts during the flare-up, considering that the reporting and investigation of live alerts was already high in the county before the flare-up.

The improvements on the indicators of triage (from 60% during the first assessment to 77% on reassessment; P=0.002) suggest that the mentoring of the staff during the first assessment led to behavioral change among the health workers, leading to better triaging for EVD when people visited the facility. The on-site training of the staff also improved the indicators pertaining to personal/staff training from 78% to 89% (P=0.04), since these indicators do not require any interventions other than on-site mentoring. The mentoring of the staff also contributed to an improvement in the system for checking and reporting staff health issues in the facilities, as well as led to the permanent presence of trained clinicians whenever the facility was open. However, no significant improvement was verified in other indicators like equipment and infrastructure for isolation units, which require interventions from higher administration levels, since the procurement of supplies and construction of infrastructure may be slow as the county is not currently reporting any EVD positive case. A similar assessment conducted in Sierra Leone in 2014 led to a quick intervention from stakeholders and partners evidenced by the immediate provision of equipments and supplies, considering that, unlike Sinoe County in Liberia, these districts were facing an active EVD outbreak (29).

The presence of trained staff in 90% of the health facilities; the small attrition of RRT members, contact tracers, and trained burial team members trained during the original outbreak; the availability of stock of IPC supplies for at least one month in 27 (90%) facilities; and other logistics, including the availability of three vehicles rendered the reactivation of the multi-sectoral EVD control strategy fast and relatively affordable. In addition, despite the lack of supplies to attain full functionality, the presence of an isolation space for receiving patients in 43% of the health facilities, and the presence of an ETU with a capacity of 60 beds and a CCC with a capacity of 12 beds suggest that few more interventions would be required to respond any eventual importation of cases from Monrovia or elsewhere.

Our article has several potential limitations. First, the assessments were conducted as emergency interventions, and not enough time was available to train the personnel who conducted these assessments. However, the personnel received orientations during a two-hour meeting that was conducted before the assessment. In addition, a new improved tool for reassessment was introduced before the end of the flare-up. Therefore, two different tools were used for assessment and reassessment, although some indicators did not change. Thus, only 17 (49%) facilities were assessed and re-assessed using the same tool. To minimize this limitation, our analysis only included the indicators present in the both tools. The time interval from assessment to reassessment was not the same for all the health facilities, and it varied from one to five months. Thus, some facilities may have had more time to improve than others did.

Additionally, there may have been an information bias due the fatigue of the interviewed staff since the facilities were receiving more visitors than usual during the flare-up. This may have led them to provide answers that would not require follow-up questions. Further, the interventions were implemented when the county did not report any confirmed EVD case. Thus, some key areas of EVD response, like access to food and other supplies by contacts during contact tracing and case management, were not assessed. Our study did not determine if the supplies provided and the staff trained will remain available in the county if a flare-up occurred two or more years after the end of the original outbreak, and the implications of the same. Despite these potential limitations, our findings may be taken into account for assessing the preparedness for EVD and other future outbreaks, leading to improved surveillance, early detection, and control, as well as prevention of infection among health workers.

In conclusion, as part of outbreak preparedness, community engagement may contribute to the report of community-based events, specifically community deaths for EVD surveillance. The mentoring of the staff at health facilities, combined with the assessment of IPC, would lead to behavioral change among the health workers, thereby increasing the IPC compliance and improving outbreak surveillance. The low attrition among the personnel trained in outbreak response, and presence of supplies at health facilities made easier, faster, and affordable to achieve the reactivation of the response structures.

We recommend a periodic reassessment of IPC supplies and equipment in health facilities, combined with mentoring of health workers, early advocacy for partners and stakeholders to provide the required equipment and to facilitate the construction of isolation units, and the implementation of reinforcement measures to reduce attrition among the trained health workers, especially within the first year after the end of any outbreak.

## Acknowledgements

We thank Benjamim Karmo, Leleh Gornorpewu, Anthony Moore and Matirankie Kanneh for the support provided for the data analysis, social mobilization and assessments conducted in Sinoe County during the flare-up in Monrovia.

We would also like to show our gratitude to Medical Teams International for orientations provided to the rapid response team in Sinoe County and United Nations International Children’s Emergency Fund for the technical support provided for social mobilization activities.

## Data availability

All data supporting this study are openly available from figshare, DOI 10.6084/m9.figshare.4902929 at https://figshare.com/s/079e2e55fc7b3948993d.

